# PTEN is required for human Treg suppression of costimulation

**DOI:** 10.1101/2022.03.06.483188

**Authors:** Avery J. Lam, Manjurul Haque, Kirsten A. Ward-Hartstonge, Prakruti Uday, Christine M. Wardell, Jana K. Gillies, Madeleine Speck, Majid Mojibian, Ramon I. Klein Geltink, Megan K. Levings

**Affiliations:** BC Children’s Hospital Research Institute, Vancouver, BC, V5Z 4H4, Canada; Department of Surgery, University of British Columbia, Vancouver, BC, V5Z 1M9, Canada; Department of Pathology and Laboratory Medicine, University of British Columbia, Vancouver, BC, V6T 2B5, Canada; Department of Molecular Oncology, BC Cancer Research, Vancouver, BC, V5Z 1L3, Canada; School of Biomedical Engineering, University of British Columbia, Vancouver, V6T 1Z3, Canada; Department of Pathology, Stanford University, Stanford, CA, 94305, USA; Department of Microbiology and Immunology, University of Otago, Dunedin, 9016, New Zealand

**Keywords:** CRISPR-Cas9, immune regulation, PI3K-AKT, PTEN, regulatory T cells

## Abstract

Regulatory T cell (Treg) therapy is under clinical investigation for the treatment of transplant rejection, autoimmune disease, and graft-versus-host disease. With the advent of genome editing, attention has turned to reinforcing Treg function for therapeutic benefit. A hallmark of Tregs is dampened activation of PI3K-AKT signalling, of which PTEN is a major negative regulator. Loss-of-function studies of PTEN, however, have not conclusively shown a requirement for PTEN in upholding Treg function and stability. Using CRISPR-based genome editing in human Tregs, we show that PTEN ablation does not cause a global defect in Treg function and stability; rather, it selectively blocks their ability to suppress antigen-presenting cells. PTEN-KO Tregs exhibit elevated glycolytic activity, upregulate FOXP3, maintain a Treg phenotype, and have no discernable defects in lineage stability. Functionally, PTEN is dispensable for human Treg-mediated inhibition of T cell activity in vitro and in vivo but is required for suppression of costimulatory molecule expression by antigen-presenting cells. These data are the first to define a role for a signalling pathway in controlling a subset of human Treg activity. Moreover, they point to the functional necessity of PTEN-regulated PI3K-AKT activity for optimal human Treg function.

## Introduction

Regulatory T cells (Tregs) suppress unwanted immune responses. Cell therapy with Tregs is currently under clinical investigation for the prevention or treatment of conditions of overactive immunity, including transplant rejection, autoimmune disease, and graft-versus-host disease. The advent of genome editing with CRISPR/Cas9 opens a new frontier for developing next-generation Treg cell therapies with tailored functional profiles, specificities, and enhanced stability (Amini et al., 2021; Ferreira et al., 2019). These tools also enable interrogation of pathways governing Treg biology and the consequent rational design of therapeutic, gene-edited Tregs.

TCR engagement, costimulation, and IL-2 signals engage the PI3K-AKT pathway to enable T cell proliferation, metabolism, and function (Fan and Turka, 2018). During PI3K-AKT activation, PI3K phosphorylates the membrane phospholipid PIP2 generating PIP3, which acts as an anchor for AKT. AKT can then be phosphorylated at S473 and T308 by the kinases mTORC2 and PDK1, respectively; phosphorylation at both sites is required for full activation of AKT (Manning and Toker, 2017). A major target downstream of PI3K-AKT is mTORC1, which among many other targets ultimately phosphorylates ribosomal protein S6 and initiates ribosomal protein synthesis. PTEN opposes PI3K-AKT activation by catalyzing the reverse reaction, dephosphorylating PIP3 into PIP2 (Manning and Toker, 2017).

A hallmark of Tregs is altered PI3K-AKT signalling compared to CD4^+^CD25^−^ conventional T cells (Tconvs). Downstream of TCR, CD28, or IL-2 signals, Tregs exhibit comparable PI3K activity (Bensinger et al., 2004) but dampened AKT activation, particularly at the mTORC2-dependent S473 phosphorylation site (Crellin et al., 2007). Low AKT activity is functionally relevant in Tregs, as overexpression of constitutively active AKT inhibits mouse Treg development in vivo and human Treg suppressive function in vitro (Crellin et al., 2007; Haxhinasto et al., 2008). This effect on Treg dysfunction is associated with increased activation of the AKT target mTORC1: functionally deleterious AKT overactivation in Tregs causes increased p-S6 (Crellin et al., 2007), and the mTORC1 inhibitor rapamycin potentiates de novo Foxp3 induction in Tconvs (Haxhinasto et al., 2008; Sauer et al., 2008). Reduced AKT activation is also important for maintaining Foxo transcription factor activity, which is needed for Treg development and function (Ouyang et al., 2012). Foxo1 and Foxo3 promote Foxp3 expression, upregulate CTLA-4 and other Treg-associated genes, and inhibit expression of the inflammatory cytokine IFN-γ (Kerdiles et al., 2010; Ouyang et al., 2010, 2012). On the other hand, some degree of PI3K and mTORC1 activity is required for Treg development and function (Patton et al., 2006; Zeng et al., 2013) and a Foxo1-dependent transcriptional program is needed for Treg differentiation into an activated phenotype (Luo et al., 2016). Overall, strategies to modulate the PI3K-AKT pathway may reinforce Treg function for therapeutic benefit.

PTEN loss-of-function studies in Tregs have yielded conflicting findings. Evidence supporting a beneficial role of PTEN include the finding that mice with a Treg-specific deletion of *Pten* develop an autoimmune lymphoproliferative disease characterized by excessive Th1, Tfh, and B cell responses (Huynh et al., 2015; Shrestha et al., 2015). Tregs in these mice sequentially downregulate CD25 and Foxp3 and remethylate the Treg-specific methylation region (TSDR) (Huynh et al., 2015; Shrestha et al., 2015), suggesting a loss of Treg lineage stability (Floess et al., 2007). PTEN-deficient Tregs also exhibit elevated glycolysis and increased phosphorylation of AKT at S473, indicative of mTORC2 activity (Huynh et al., 2015; Shrestha et al., 2015). Concomitant deletion of mTORC2 in Tregs rescues mice from disease and reverses the aberrant Treg phenotype (Shrestha et al., 2015). In a less penetrant strain of Treg-specific *Pten* ablation, mice do not develop autoimmunity until later in life, although their Tregs are unable to suppress anti-tumour responses (Sharma et al., 2015). Analogously in human Tregs, pharmacological inhibition of PTEN or knockdown of the PTEN-recruiting scaffold protein DLGH1 reduces their suppressive function in vitro (Kitz et al., 2016; Zanin-Zhorov et al., 2012); PTEN inhibitor-treated Tregs also upregulate IFN-γ expression (Kitz et al., 2016). On the other hand, PTEN-deficient Tregs from mice with a pan-T cell-specific deletion of *Pten* retain their suppressive function both in vitro and in an adoptive transfer model of colitis (Walsh et al., 2006). Moreover, patients with deleterious heterozygous mutations in *PTEN* exhibit normal Treg frequencies in blood and lymphoid tissue and unchanged Treg expression of CD25 and FOXP3 (Chen et al., 2017). Thus, how PTEN controls human Treg function remains ambiguous.

Seeking to understand how human Treg function and stability is controlled by PTEN, we used CRISPR/Cas9 to genetically ablate *PTEN* in human Tregs. We interrogated how loss of PTEN affected the PI3K-AKT-FOXO signalling axis in Tregs as well as their stability, phenotype, metabolic activity, and in vitro and in vivo suppressive function.

## Results

### Human PTEN^KO^ Tregs exhibit constitutive mTORC2-AKT activation

We recently established a human Treg expansion system amenable to CRISPR/Cas9 editing (Lam et al., 2021), in which flow-sorted Tregs are preactivated for 5 days, electroporated with a CRISPR/Cas9 ribonucleoprotein, and expanded for 7 days before resting overnight (**Fig. S1A–B**). To investigate the role of PTEN in human Treg biology, we designed guide RNAs (gRNAs) targeting 5′-proximal exons expressed in all splice isoforms of *PTEN* (**Fig. 1A**) and evaluated editing efficiencies in human naive Tregs (nTregs), 3 days or 8 days after editing. *PTEN*-targeting gRNAs exhibited a wide range (0–92%) of indel formation, with percent indel formation higher at 8 days post-editing compared to 3 days post-editing (**Fig. 1B**). As CRISPR/Cas9 ribonucleoproteins cleave their target DNA within hours and are degraded rapidly in cells (Kim et al., 2014), the change in indels over time suggests competitive outgrowth of PTEN-targeted nTregs in culture. Indeed, the highest-performing gRNAs by indel formation (CR4, CR5, CR9) yielded the greatest increase in fold expansion (**Fig. 1C**). The gRNAs CR5 and CR9 also resulted in efficient ablation of PTEN protein expression, 8 days after editing (**Fig. 1D**). Collectively, these findings are consistent with the anti-proliferative effects of PTEN on mouse Tregs in vitro and in vivo (Huynh et al., 2015; Shrestha et al., 2015; Walsh et al., 2006).

**Fig. 1.**
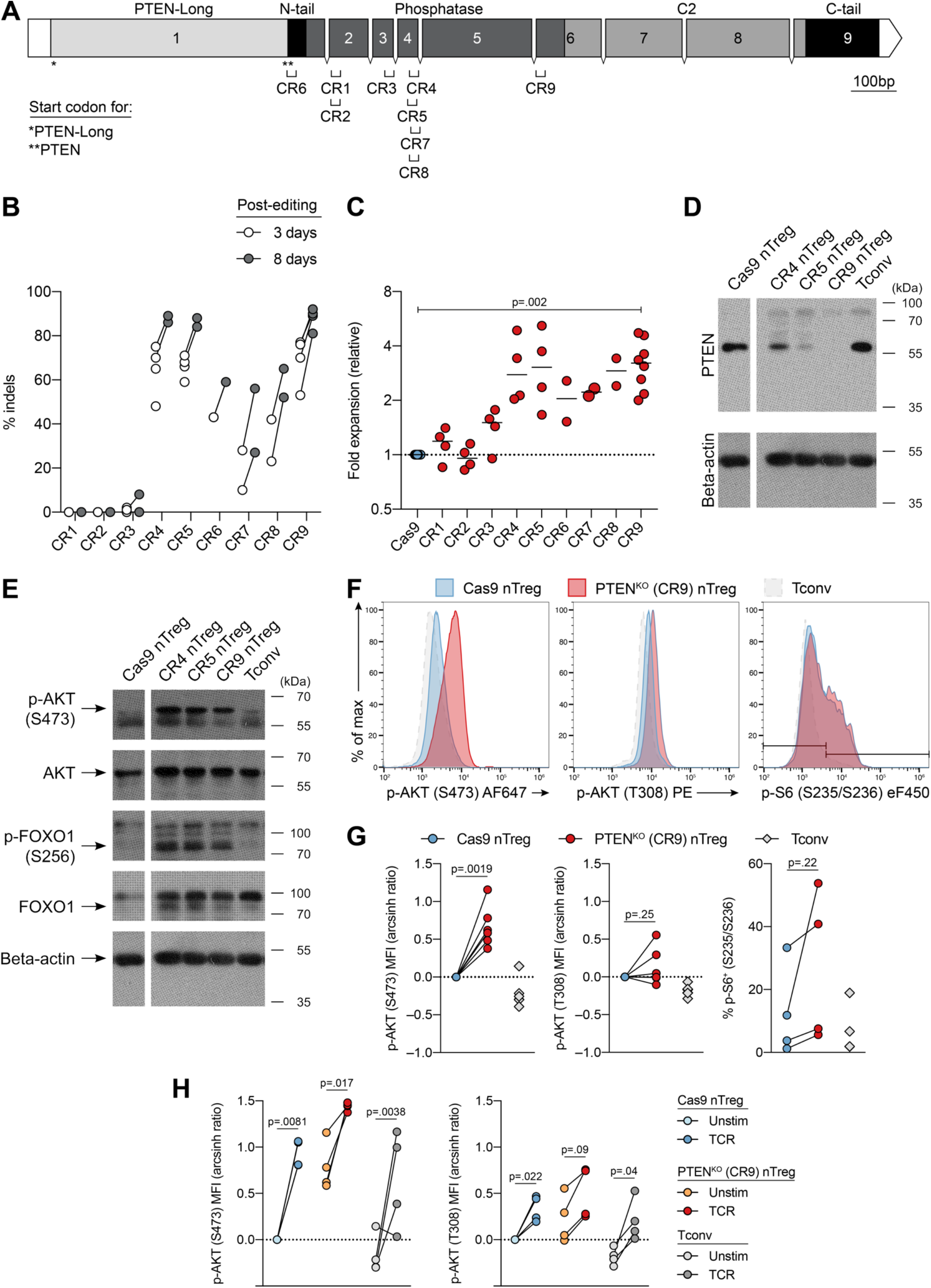
Human PTEN^KO^ Tregs exhibit constitutive mTORC2-AKT activation. (**A**) *PTEN* gene diagram with exons (1–9), introns, protein domains, and gRNA (CR1–CR9) target locations. Start codons of *PTEN* gene products indicated with * (PTEN-Long) or ** (PTEN). 5′UTR, introns, and 3′UTR not to scale. (**B–E**) nTregs were edited with the indicated *PTEN*-targeting gRNA. (**B**) Percent indel formation, 3 days and 8 days post-editing (n=2–4, 1–2 experiments). (**C**) Fold expansion within days 0–12, relative to Cas9 nTregs (n=2–10, 1–5 experiments). (**D**) PTEN expression by western blotting, 8 days post-editing (n=2, 1 experiment). (**E**) Expression of p-AKT (S73), AKT, p-FOXO1 (S256), and FOXO1 by western blotting, 8 days post-editing (n=2, 1 experiment). (**F–H**) nTregs were edited with PTEN CR9 gRNA (13 days), then starved overnight without serum or IL-2. Example (**F**) and quantified (**G**) expression of p-AKT (S473), p-AKT (T308), and p-S6 (S235/S236) by flow cytometry without any stimulation. (**H**) Cells were TCR-activated for 5 min. p-AKT (S473) and p-AKT (T308) expression by flow cytometry. Dots in (**B–C**), (**G–H**) represent individual donors. Solid horizontal lines in (**C**) represent medians. In (**D–E**), lanes from the same blot have been reordered for clarity, and indicated with a white vertical gap. Significance determined by mixed-effects model with Geisser-Greenhouse correction and Dunnett’s multiple comparisons test (all groups compared to Cas9) in (**C**), paired t-test in (**G**), and matched 2-way ANOVA with Sidak’s multiple comparisons test in (**H**). Tconvs shown for reference. MFI, geometric mean fluorescence intensity.

As PTEN inhibits the PI3K-AKT pathway, we tested the effects of PTEN ablation on this pathway in nTregs. After overnight rest with reduced IL-2, Cas9 nTregs exhibited minimal phosphorylation at AKT (S473) or its substrate FOXO1 (S256) by phospho-western blotting (**Fig. 1E**). In contrast, nTregs edited with the *PTEN*-targeting gRNAs CR4, CR5, or CR9 exhibited greater basal p-AKT (S473) and p-FOXO1 (S256) expression (**Fig. 1E**), indicative of AKT activation and FOXO1 inactivation. We selected CR9 to target PTEN in nTregs for further characterization due to its efficient genomic editing, protein ablation, and expected effects on AKT activity.

nTregs edited with the PTEN-targeting gRNA CR9 (hereafter PTEN^KO^ nTregs) were further starved overnight without any IL-2 or serum for phospho-flow cytometry. PTEN^KO^ nTregs consistently exhibited increased basal phosphorylation of AKT at S473, but not at T308 or of ribosomal protein S6 (S235/S236) (**Fig. 1F–G**). We also asked whether PTEN^KO^ nTregs had altered TCR-mediated AKT activation. After brief (5 min) activation with soluble anti-CD3 and anti-CD28, Cas9 nTregs increased phosphorylation of AKT at both S473 and T308, as expected (**Fig. 1H**). TCR activation further increased p-AKT (S473) in PTEN^KO^ nTregs beyond elevated basal levels. Meanwhile, p-AKT (T308) increased comparably in Cas9 and PTEN^KO^ nTregs upon TCR activation (**Fig. 1H**). Overall, as with mice (Huynh et al., 2015; Shrestha et al., 2015), PTEN in human Tregs preferentially inhibits mTORC2-AKT signalling.

### Human PTEN^KO^ Tregs have elevated glycolytic activity

Human Tregs have a distinct metabolic profile relative to Tconvs: ex vivo Tregs are highly glycolytic, while in vitro proliferating Tregs use both glycolysis and fatty acid oxidation (Procaccini et al., 2016). Analogously, mouse Tregs require glycolysis for their proliferation and suppressive capacity (Gerriets et al., 2016; Zeng et al., 2013). Unrestrained glycolysis, however, such as through increased PI3K-AKT activation, may be detrimental to mouse and human Treg function (De Rosa et al., 2015; Gerriets et al., 2016; Saravia et al., 2020). We therefore asked whether human PTEN^KO^ nTregs exhibited altered metabolic activity.

After expansion, we performed Seahorse extracellular flux analysis on Tregs without any further stimulation, upon the sequential addition of oligomycin (inhibits mitochondrial ATP synthase), FCCP (uncouples mitochondria), and a combination of rotenone and antimycin A (inhibits mitochondrial complexes I and III) (**Fig. 2A–B**). Basal glycolysis and glycolytic capacity were determined by the extracellular acidification rate (ECAR) before and after the addition of oligomycin (which maximizes the glycolytic rate), respectively (**Fig. 2A**). In parallel, the oxygen consumption rate (OCR) was measured to determine basal, ATP-linked (after oligomycin), or reserve capacity of mitochondrial respiration (after FCCP) (**Fig. 2B**). PTEN^KO^ nTregs exhibited both higher basal glycolysis and higher glycolytic capacity than Cas9 nTregs, as determined by basal and maximal ECAR (**Fig. 2A, Fig. 2C**). In contrast, Cas9 and PTEN^KO^ nTregs exhibited similar ATP-linked mitochondrial respiration and reserve respiratory capacity (**Fig. 2B, Fig. 2D**), suggesting similar mitochondrial fitness. As a result, PTEN^KO^ Tregs had a significantly lower OCR:ECAR ratio, a proxy for relative metabolic activity, compared to Cas9 controls activity (**Fig. 2E**). These data confirm the expected effect of PTEN^KO^ on Tregs in terms of preferential use of glycolysis, as with their mouse counterparts (Huynh et al., 2015; Shrestha et al., 2015).

**Fig. 2.**
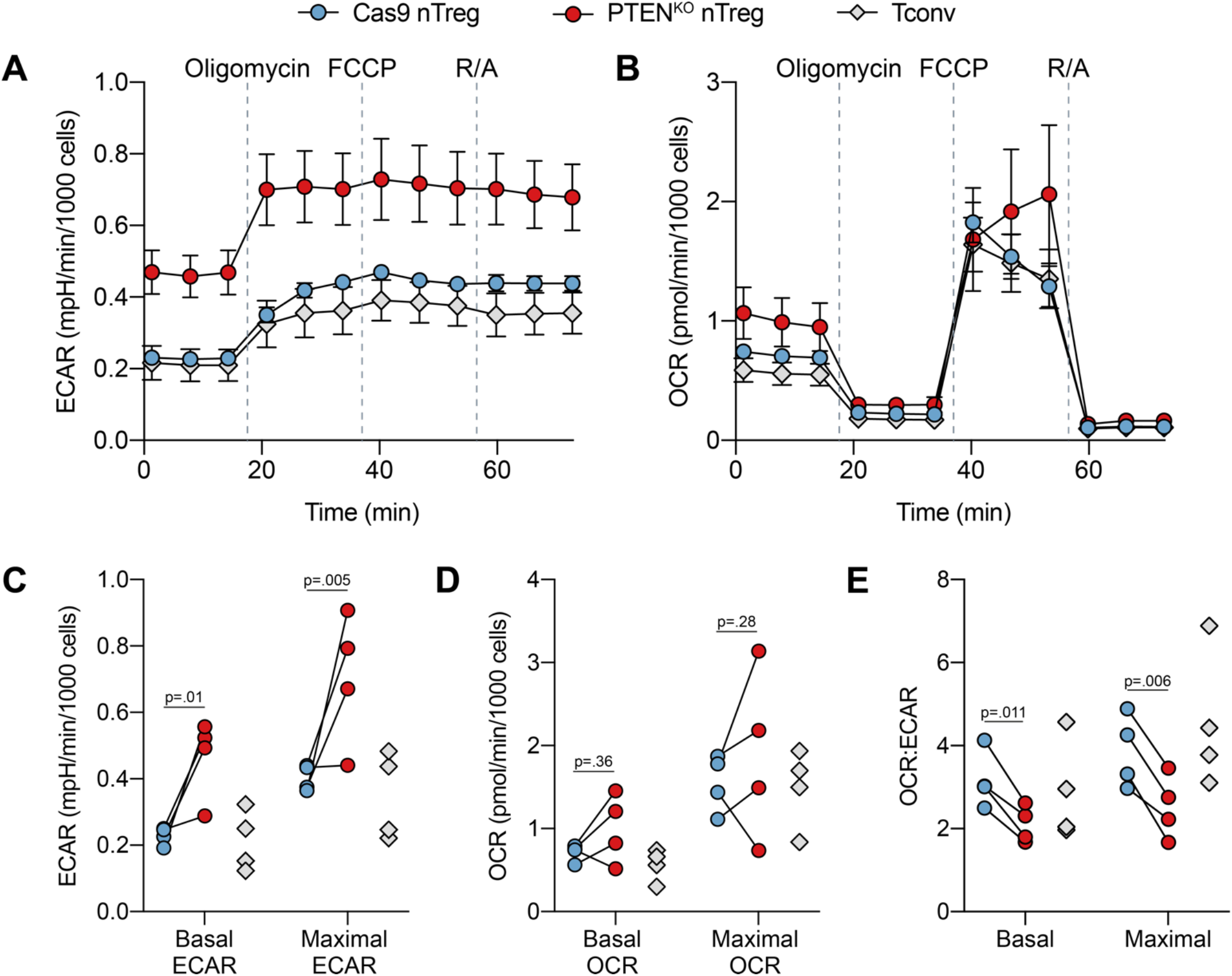
Human PTEN^KO^ Tregs have elevated glycolytic activity. (**A–E**) Expanded Cas9 nTregs, PTEN^KO^ nTregs, and Tconvs (12 days) were sequentially treated with oligomycin, FCCP, and rotenone + antimycin A (R/A) as indicated, in the Seahorse Mito Stress Test (n=4, 2 experiments). (**A**) Extracellular acidification rate (ECAR) and (**B**) oxygen consumption rate (OCR) over time. (**C–E**) Basal and maximal (**C**) ECAR, (**D**) OCR, and (**E**) OCR:ECAR ratio. Symbols in (**A–B**) represent mean±SEM. Dots in (**C–E**) represent individual donors. Each measurement is the average of three technical replicates. Significance in (**C–E**) determined by matched 2-way ANOVA with Sidak’s multiple comparisons test. Tconvs shown for reference.

### PTEN^KO^ Tregs have preserved lineage stability

In mice, PTEN^KO^ Tregs have impaired lineage stability and sequentially lose CD25 and Foxp3 expression over time both in vitro and in vivo (Huynh et al., 2015; Shrestha et al., 2015). We were thus surprised to find that human PTEN^KO^ nTregs had consistently higher FOXP3 expression compared to their Cas9 counterparts, with no change in CD25 expression (**Fig. 3A–B**). Expression of other characteristic Treg proteins, including the coinhibitory molecule CTLA-4 and transcription factor Helios, were also unchanged upon PTEN ablation (**Fig. 3B**). Consistent with maintenance of a stable Treg phenotype, there was sustained demethylation of the *FOXP3* TSDR in PTEN^KO^ nTregs (**Fig. 3C**).

**Fig. 3.**
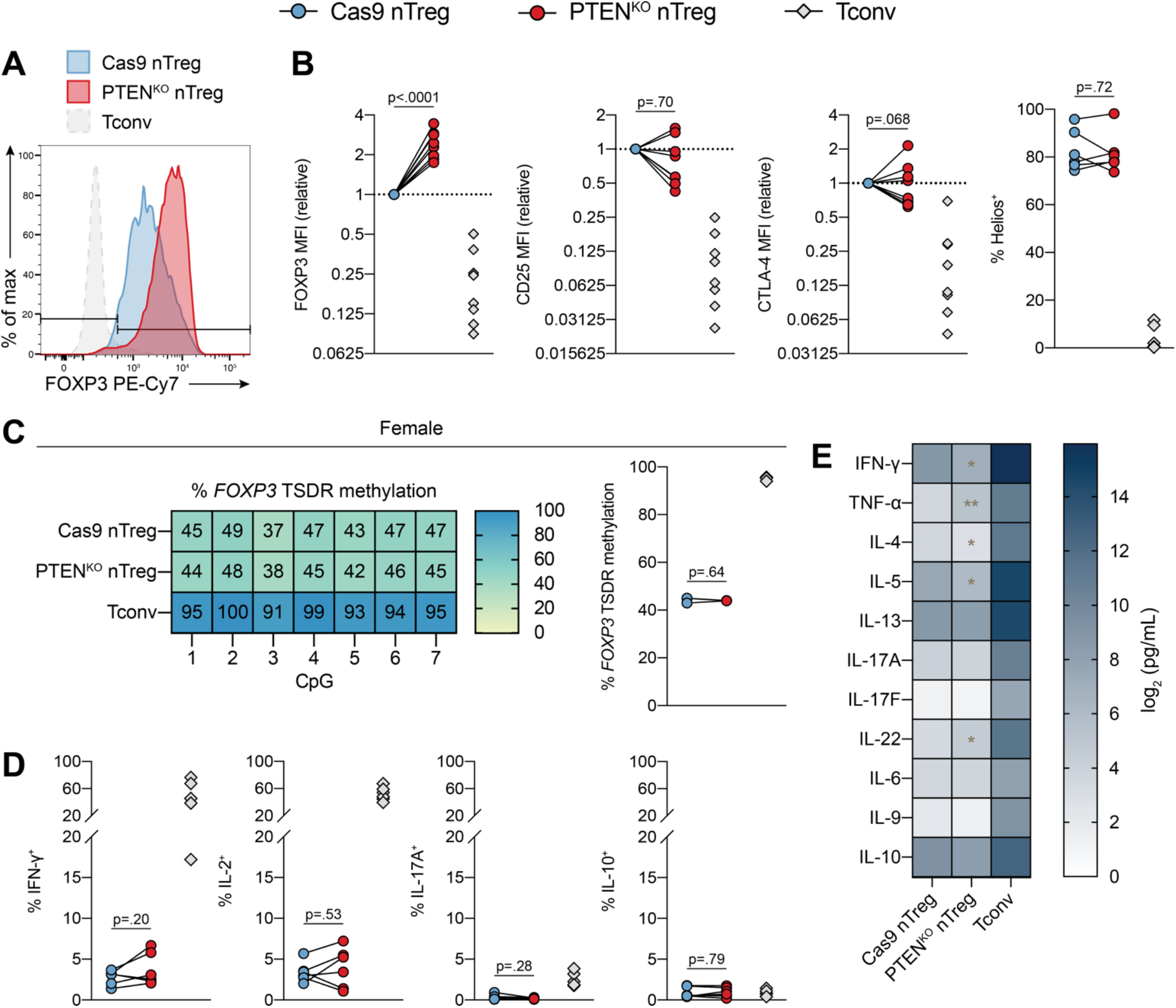
Human PTEN^KO^ Tregs have preserved lineage stability. (**A–F**) Cas9 nTregs, PTEN^KO^ nTregs, and Tconvs were expanded for total 13 days. (**A**) Example FOXP3 expression. (**B**) FOXP3, CD25, CTLA-4, and Helios expression by flow cytometry (n=8, 4 experiments). (**C**) Example (left) and average (right) *FOXP3* TSDR CpG methylation analysis (n=3 females, 2 experiments). (**D**) IFN-γ, IL-2, IL-17A, and IL-10 expression by flow cytometry after restimulation with PMA, ionomycin, and brefeldin A (6 h) (n=6, 3 experiments). (**E**) Secretion of the indicated cytokines into culture supernatants after activation with IL-2 and TCR (4 days) (n=8, 4 experiments). Dots in (**B–D**) represent individual donors. (**E**) depicts means. Significance in (**B–E**) determined by paired t-test of PTEN^KO^ vs. Cas9 nTreg; * (p < 0.05) or ** (p < 0.01) indicate a significant difference from Cas9 nTreg. Tconvs shown for reference. MFI, geometric mean fluorescence intensity.

A hallmark of Tregs is low production of inflammatory cytokines relative to Tconvs; reversal of this phenotype is associated with lineage instability. To further explore the role of PTEN in Treg stability, we measured the effect of PTEN ablation on cytokine production by intracellular staining or cytometric bead array. Cas9 and PTEN^KO^ nTregs were equivalent in terms of low intracellular expression of IFN-γ, IL-2, IL-17A, and IL-10 (**Fig. 3D**). For cytokines secreted into the culture supernatant, PTEN^KO^ nTregs had a modest downregulation of secretion of IFN-γ, IL-4, and IL-5 relative to Cas9 nTregs and upregulation of TNF-α and IL-22 (**Fig. 3E**). Collectively, these data suggest that loss of PTEN in nTregs does not result in lineage instability.

### Superior suppression of T cell proliferation by human PTEN^KO^ Tregs

The extent to which PTEN influences Treg function is unclear. Depending on the context and model, mouse PTEN-deficient Tregs are equally, partially, or not able to suppress T cell proliferation in vitro (Delgoffe et al., 2013; Sharma et al., 2015; Walsh et al., 2006) and T cell responses in vivo (Sharma et al., 2015; Shrestha et al., 2015; Walsh et al., 2006). In human cells, Tregs pre-treated with a pharmacological inhibitor of PTEN are less able to suppress T cell proliferation in vitro (Kitz et al., 2016). To comprehensively determine the role of PTEN in human Tregs, we evaluated the suppressive capacity of PTEN^KO^ nTregs in three different contexts: suppression of T cell proliferation, antigen-presenting cell (APC) function, and xenogeneic graft-versus-host disease (GVHD).

To assess Treg-mediated suppression of T cell activity in vitro, Tregs were co-cultured with purified CD3^+^ T cells and anti-CD3/anti-CD28 beads for 4 days. In this assay, Tregs directly suppress T cell proliferation and cytokine secretion in an APC-independent manner. We found that PTEN^KO^ nTregs were substantially better able to suppress the proliferation of both CD4^+^ and CD8^+^ T cells than Cas9 nTregs, particularly at suboptimal Treg ratios (**Fig. 4A**). Similarly, PTEN^KO^ Tregs more potently suppressed T cell secretion of the Th1 cytokines IFN-γ and IL-2, though not of the Th17 cytokines IL-17A or IL-22 (**Fig. 4B**). Superior T cell-suppressive function of gene-edited nTregs persisted 15 days after editing (**Fig. S2A**). A similar effect was seen in memory Tregs (mTregs): PTEN^KO^ mTregs exhibited a more potent suppressive effect on CD4^+^ and CD8^+^ T cell proliferation, 15 days after editing (**Fig. S2B**).

**Fig. 4.**
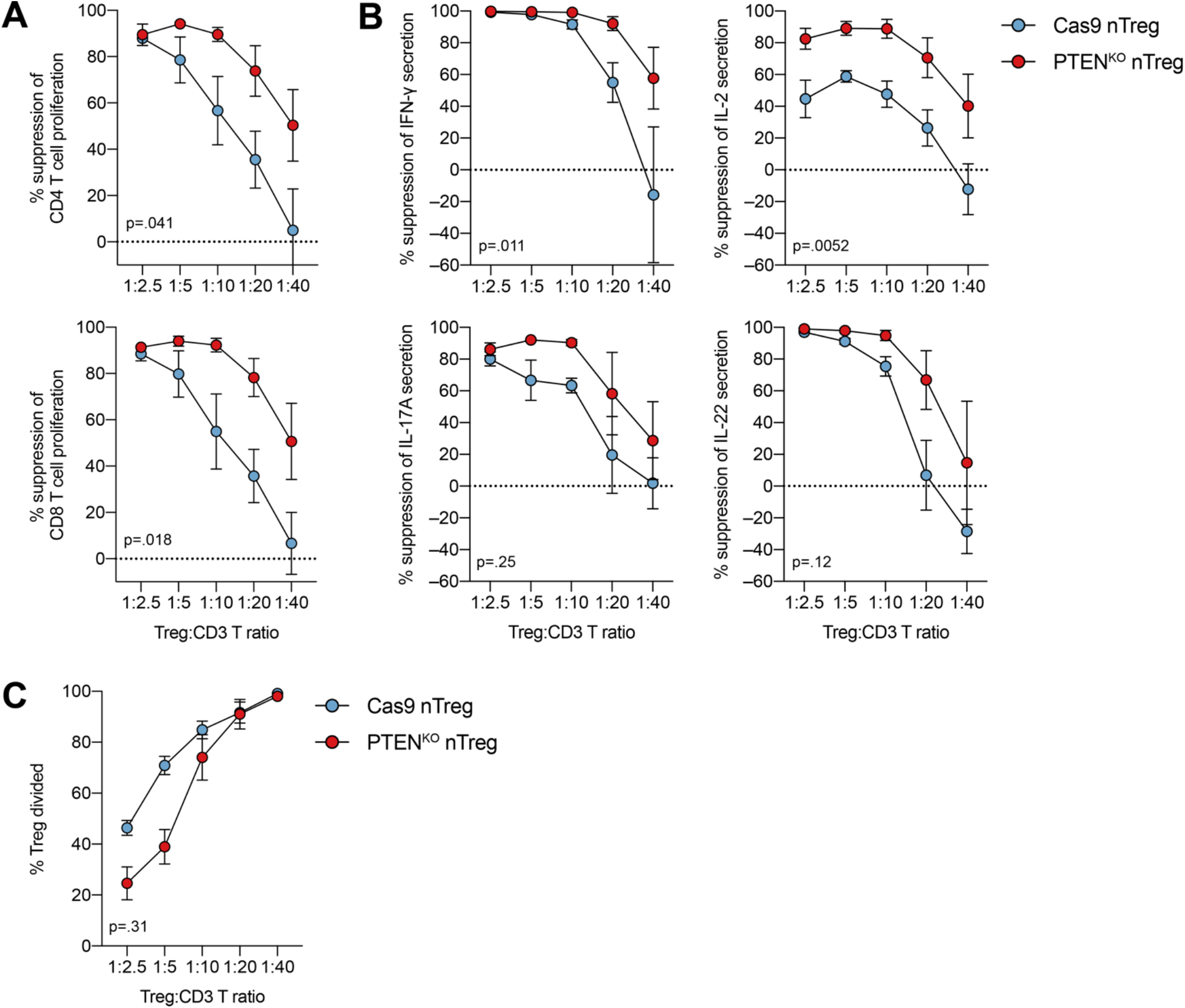
Superior suppression of T cell proliferation by human PTEN^KO^ Tregs. (**A–C**) Cas9 and PTEN^KO^ nTregs were expanded for total 13 days. (**A**) Tregs and allogeneic CD3^+^ T cells were co-cultured at the indicated ratios and TCR-activated (4 days). (**A**) Suppression of CD4^+^ T cell (left) and CD8^+^ T cell (right) proliferation (n=4, 2 experiments). (**B**) Suppression of IFN-γ, IL-2, IL-17A, and IL-22 secretion (n=4, 2 experiments). (**C**) Proliferation of Tregs in the Treg:CD3^+^ T cell co-culture (n=6, 3 experiments). (**A–C**) depict mean±SEM. Significance in (**A–C**) determined by t-test of the areas under the curve.

Given that PTEN^KO^ nTregs can outgrow their unedited Treg counterparts within a mixed population (**Fig. 1B–C**), we considered the possibility that the increased suppression was related to PTEN^KO^ nTreg-mediated nutrient depletion. This was unlikely the case, however, as PTEN^KO^ nTregs proliferated to a similar extent to Cas9 nTregs during co-culture with allogeneic CD3^+^ T cells (**Fig. 4C, Fig. S2C**). Thus, in an APC-free in vitro suppression assay, PTEN ablation increases the ability of human nTregs and mTregs to suppress T cell proliferation and secretion of Th1 cytokines.

### Suppression of xenogeneic graft-versus-host-disease by human PTEN^KO^ Tregs

We next evaluated human PTEN^KO^ Treg function in vivo with a xenogeneic model of GVHD, wherein polyclonal Tregs control GVHD induced by human PBMCs in immunodeficient NSG mice (Mutis et al., 2006). Mice were injected with PBMCs in the presence or absence of Cas9 or PTEN^KO^ nTregs (**Fig. 5A, Fig. S3A**). We employed high (1:2) and low (1:4) Treg:PBMC ratios (**Fig. 5A**), since we previously found that defects in human Treg function in vivo were more apparent at lower Treg:PBMC ratios (Dawson et al., 2020). To distinguish PBMCs and Tregs, we used differential HLA-A2 expression with HLA-A2^−^ PBMCs and HLA-A2^+^ Tregs. As expected, mice receiving PBMCs alone rapidly developed GVHD, with all mice reaching a humane endpoint within 28 days (**Fig. 5B**). Co-injection of Cas9 or PTEN^KO^ nTregs at either a high or low ratio significantly delayed the onset of GVHD (**Fig. 5B**). The survival distributions of mice receiving PTEN^KO^ nTregs at low or high ratios were not significantly different compared to mice receiving Cas9 nTregs (high ratio). Consistent with survival data, mice in PTEN^KO^ nTreg-containing groups developed a combined GVHD score (based on GVHD symptoms and weight loss) at similar rates to those receiving Cas9 nTregs (**Fig. 5C**).

**Fig. 5.**
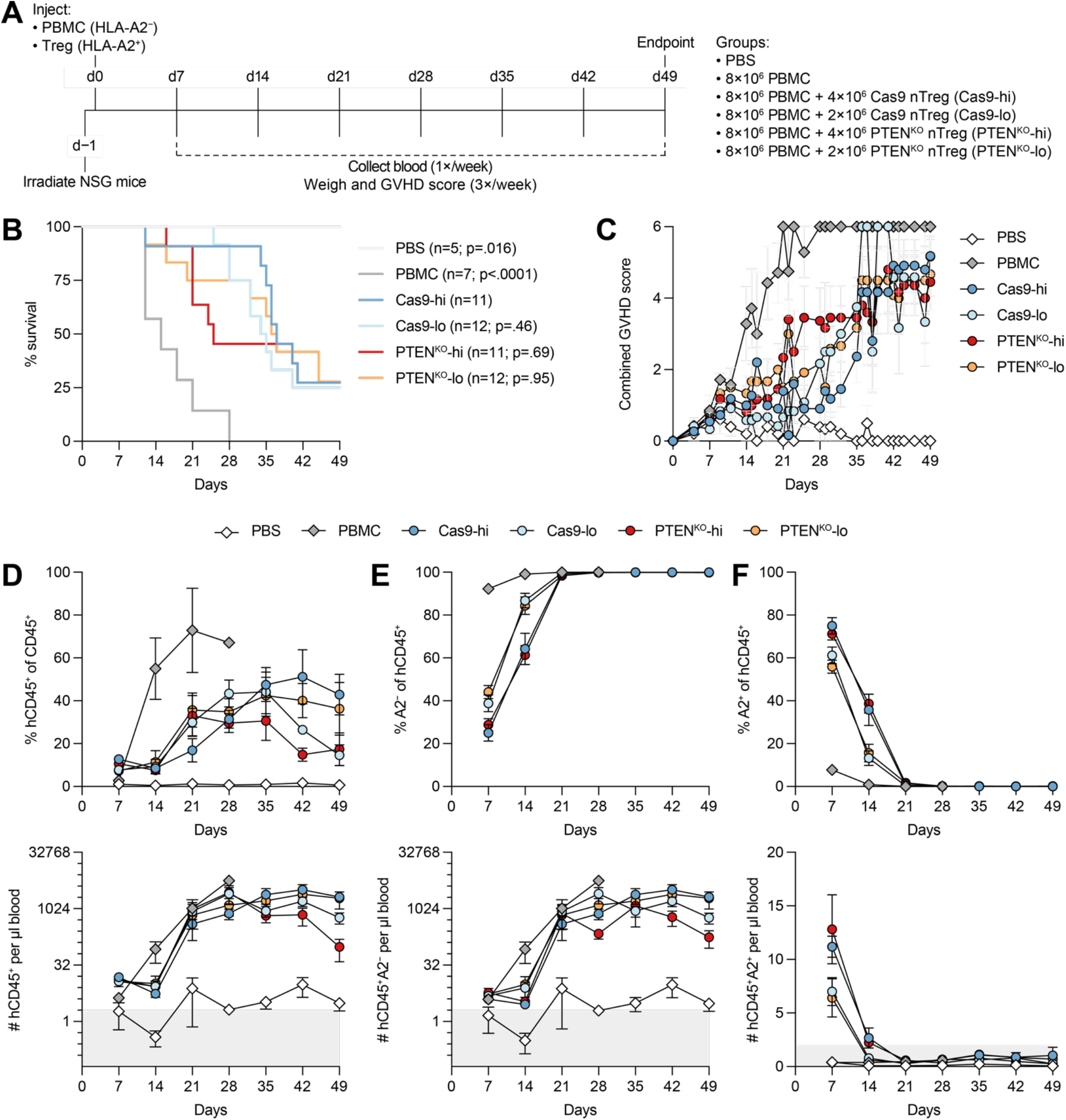
Suppression of xenogeneic graft-versus-host-disease by human PTEN^KO^ Tregs. (**A**) NSG mice were irradiated 1 day before injection of PBS or PBMCs (8 × 10^6^/mouse) in the presence or absence of the indicated nTregs (4 × 10^6^/mouse (hi) or 2 × 10^6^/mouse (lo)) (n=2 PBS mice, n=4 PBMC mice, n=6 Cas-hi mice, n=6 Cas-lo mice, n=5 PTEN-hi mice, n=5 PTEN-lo mice; pooled from 2 cohorts). Differential HLA-A2 expression (HLA-A2^−^ PBMCs, HLA-A2^+^ nTregs) was used to distinguish the transferred populations. Mice were scored for GVHD thrice weekly and bled once weekly. (**B**) Survival curves. (**C**) Combined GVHD scores over time. (**D–F**) Proportion (top) and number (bottom) over time of (**D**) hCD45^+^ cells within total mouse/human CD45^+^ cells, (**E**) HLA-A2^−^ cells within hCD45^+^ cells, and (**F**) HLA-A2^+^ cells within hCD45^+^ cells. n values indicated in (**B**) (2 experiments). (**C–F**) depict mean±SEM. Significance in (**B**) determined by log-rank test (all groups compared to Cas9-hi).

We also examined the engraftment of PBMCs (HLA-A2^−^) and Tregs (HLA-A2^+^) over time. PBMC-alone mice experienced more rapid engraftment of PBMCs by 14 days after injection, whereas PBMC engraftment in all Treg-containing groups was delayed (**Fig. 5D–E**). The kinetics and extent of PBMC engraftment were not substantially different between Cas9 and PTEN^KO^ nTregs or between high and low Treg ratios (**Fig. 5E**). Consistent with previous studies (Dawson et al., 2019, 2020; MacDonald et al., 2016), Tregs comprised the majority of engrafted human CD45^+^ cells 7 days after injection but rapidly diminished by 14 days (**Fig. 5F**).

Finally, we examined the phenotype of injected Tregs 7 days after injection, at peak Treg engraftment. We found that although FOXP3 expression in Cas9 and PTEN^KO^ nTregs in vivo was comparable, PTEN^KO^ nTregs at both high and low ratios showed decreased co-expression of FOXP3 and Helios compared to Cas9 nTregs (high ratio) (**Fig. S3B**). However, decreased Treg co-expression of FOXP3 and Helios at 7 days post-injection did not correlate with shorter individual mouse survival time (**Fig. S3C**). Overall, polyclonal PTEN^KO^ nTregs are comparable to their Cas9 nTreg counterparts at delaying the onset of xenogeneic GVHD in vivo.

### Impaired suppression of dendritic cells by human PTEN^KO^ Tregs

The finding that PTEN^KO^ nTregs were more suppressive against T cells than Cas9 nTregs in vitro but not in vivo prompted us to examine their effects on other cells. We previously reported that human Treg suppression of monocyte-derived dendritic cell (moDC) costimulation better predicts in vivo function than in vitro T cell suppression assays (Dawson et al., 2020). Moreover, how expression of PTEN might affect Treg suppression of non-T cells is not known. We therefore evaluated the ability of PTEN^KO^ nTregs to suppress human moDCs.

Because antigen-specific Tregs more potently suppress moDC expression of costimulatory molecules than polyclonal Tregs (Huang et al., 2021), we generated antigen-specific PTEN^KO^ nTregs using chimeric antigen receptor (CAR) technology (MacDonald et al., 2016) to increase assay sensitivity. Cas9 and PTEN^KO^ nTregs were transduced with a CAR specific for HLA-A2 (hereafter A2-CAR), then magnetically purified for CAR expression (**Fig. S4A**). We verified that A2-CAR Cas9 and PTEN^KO^ nTregs uniformly expressed A2-CAR, as determined by expression of the Myc epitope tag (**Fig. 6A**). As with polyclonal PTEN^KO^ nTregs (**Fig. 3**), A2-CAR PTEN^KO^ nTregs exhibited a stable Treg phenotype, upregulating FOXP3 and maintaining high expression of the Treg-associated proteins CTLA-4, CD25, and Helios relative to A2-CAR Cas9 nTregs (**Fig. 6A, Fig. S4B**). Functionally, in a polyclonal T cell suppression assay, A2-CAR PTEN^KO^ nTregs exhibited a superior ability to suppress CD4^+^ and CD8^+^ T cell proliferation compared to A2-CAR Cas9 nTregs (**Fig. S4C**), consistent with our findings in **Fig. 4A** and confirming that introduction of the A2-CAR does not alter the phenotypic and functional consequences of PTEN ablation in nTregs.

**Fig. 6.**
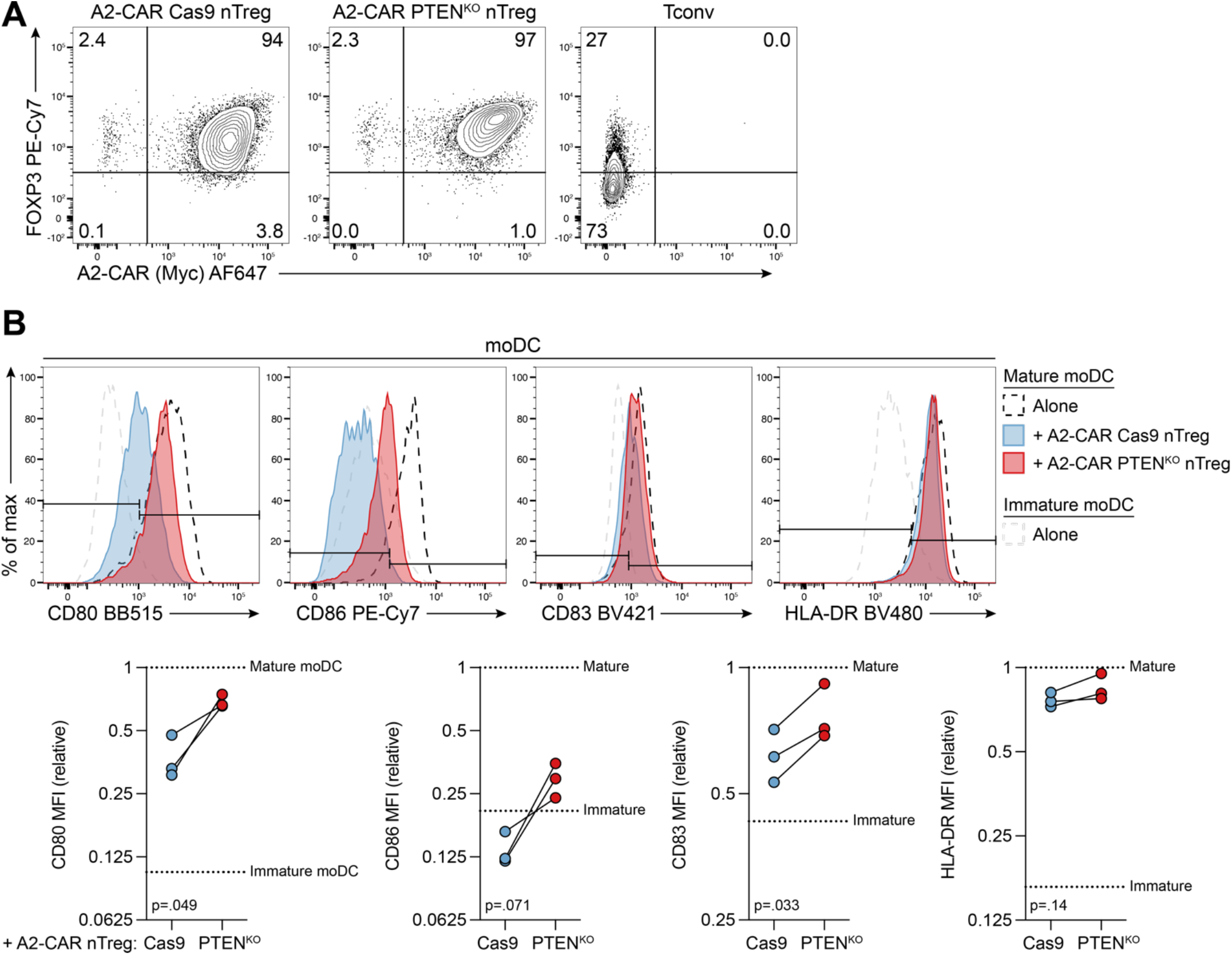
Impaired suppression of dendritic cells by human PTEN^KO^ Tregs. (**A–B**) A2-CAR Cas9 and A2-CAR PTEN^KO^ nTregs were expanded for total 13 days (n=3, 1 experiment). (**A**) Representative Treg expression of FOXP3 and myc (A2-CAR epitope tag); Tconvs shown for reference. (**B**) A2-CAR nTregs were co-cultured with mature HLA-A2^+^ moDCs (5:1) for 4 days. Example (top) and quantified (bottom) moDC expression of CD80, CD86, CD83, and HLA-DR. Dotted lines indicate immature or mature moDCs cultured alone, as indicated. Significance determined by paired t-test. MFI, geometric mean fluorescence intensity.

Next, we co-cultured A2-CAR Tregs with mature HLA-A2^+^ moDCs for 4 days (**Fig. S4A**). As expected, mature moDCs expressed high levels of the costimulatory molecules CD80, CD86, and CD83, as well as HLA-DR (**Fig. 6B**). Consistent with our previous findings, co-culture with A2-CAR Cas9 nTregs suppressed expression of CD80, CD86, and CD83, but not HLA-DR, on moDCs (**Fig. 6B**) (Dawson et al., 2020). Surprisingly, co-culture with A2-CAR PTEN^KO^ nTregs revealed a significant reduction in their inhibitory effect on moDCs. A2-CAR PTEN^KO^ nTregs resulted in significantly less suppression of CD80 and CD83, and a trend towards less suppression of CD86 (**Fig. 6B**), compared to those co-cultured with A2-CAR Cas9 nTregs. Therefore, antigen-specific PTEN^KO^ nTregs have an impaired ability to suppress the costimulatory potential of moDCs.

## Discussion

Here we used CRISPR to efficiently ablate PTEN in human Tregs to uncover its role in human Treg function and stability. PTEN^KO^ Tregs exhibited many expected characteristics of PTEN-deficient cells (Manning and Toker, 2017), including constitutive activation of the mTORC2-AKT pathway and inactivation of FOXO1. PTEN^KO^ Tregs also showed elevated glycolytic activity without any alterations in mitochondrial respiratory capacity. On the other hand, PTEN^KO^ Tregs unexpectedly retained their phenotypic and epigenetic lineage stability, and upregulated the master transcription factor FOXP3. Functionally, they exhibited variable suppressive capacity in vitro and in vivo: PTEN^KO^ Tregs were better able to suppress T cell proliferation and cytokine secretion than their Cas9 counterparts but had comparable performance in a xenogeneic model of GVHD and were less able to inhibit the costimulatory potential of dendritic cells. Thus, PTEN in human Tregs is necessary for their suppression of dendritic cells but its ablation does not cause a global defect in Treg function and stability.

Despite exhibiting similar biochemical features, human PTEN^KO^ Tregs diverge from their mouse counterparts in terms of phenotype and lineage stability. The preference of human PTEN^KO^ Tregs to activate mTORC2 over mTORC1 and favour glycolysis over oxidative phosphorylation is consistent with previous findings in mice (Huynh et al., 2015; Shrestha et al., 2015). However, human Tregs upon PTEN ablation maintained FOXP3, CD25, CTLA-4, and Helios expression, similar to patients with germline heterozygous loss-of-function *PTEN* mutations (Chen et al., 2017) but in contrast to mice.

In vivo, PTEN^KO^ Tregs modestly downregulated Helios expression, a marker of human Treg stability (Lam et al., 2022), but this did not affect FOXP3^+^ Treg numbers or significantly accelerate the onset of GHVD. Human PTEN^KO^ Tregs also retained a demethylated *FOXP3* TSDR and low production of inflammatory cytokines, distinct from mouse PTEN-deficient Tregs which lose *Foxp3* TSDR methylation and upregulate IFN-γ and IL-17A (Huynh et al., 2015; Shrestha et al., 2015). Overall, the role of PTEN in human Tregs is distinct from their mouse counterparts.

One possibility for the discrepancy in phenotype between mice and humans may be incomplete loss of PTEN function. Whereas conditional knockout mouse models have biallelic mutations, Tregs from patients with germline *PTEN* mutations have only one mutant allele (Chen et al., 2017). Although we cannot rule out the possibility that a minority of unedited or single-allele-edited Tregs conferred a dominant or cell-extrinsic effect, we achieved >80% *PTEN* mutation at the genome level and near-complete ablation of PTEN protein expression. Furthermore, in mice, even partial PTEN loss is sufficient to cause Treg loss of FOXP3 and systemic autoimmunity (Shrestha et al., 2015).

To capture the breadth of human Treg function, we used several assays to evaluate their suppressive capacity in vitro and vivo, with differing results in each setting. Tregs are thought to exert their suppressive effects primarily on APCs and T cells (Schmidt et al., 2012). In terms of effects on T cells, multiple lines of evidence support a direct effect of Tregs on CD4^+^ and CD8^+^ T cells. In lymphoid tissues, Tregs cluster around rare, IL-2-producing Tconvs (Liu et al., 2015); in target tissues, Tregs persistently interact with Tconvs and CD8^+^ effector T cells independently of dendritic cells (Miska et al., 2014). In the APC-free in vitro T cell suppression assay, it is thought that Tregs rapidly suppress multiple signalling pathways downstream of TCR engagement in Tconvs and hence proliferation and cytokine production (Chellappa et al., 2016; Schmidt et al., 2011), likely in a manner independent of CTLA-4 (Cossarizza et al., 2021; Hou et al., 2019). The specific pathways mediating this in vitro suppression remain largely unclear. With mouse cells, there is evidence for a role of CD25 in mediating IL-2 depletion (Amado et al., 2013; Chinen et al., 2016), but we did not detect differential CD25 expression on human PTEN^KO^ Tregs. Instead, PTEN ablation may be affecting other pathways involved in Treg-mediated suppression of APCs, such as expression of latent TGF-β and GARP (Cuende et al., 2015; Stockis et al., 2017) or activity of the ATP/AMP-degrading ATP enzymes CD39 and CD73 (Borsellino et al., 2007; Kobie et al., 2006).

Meanwhile, antigen-specific PTEN^KO^ Tregs exhibited defective inhibition of APC costimulation, implicating a role for PTEN in controlling CTLA-4-mediated suppression. APCs are important targets of Treg function in homeostasis: in the lymph nodes, mouse Tregs interact with dendritic cells to prevent autoreactive T cell priming (Tadokoro et al., 2006; Tang et al., 2006), likely mediated by Treg constitutive expression of CTLA-4 (Matheu et al., 2015). CTLA-4 not only outcompetes CD28 for binding to CD80 and CD86 but also captures these molecules from the surface of dendritic cells for endosomal degradation (Ovcinnikovs et al., 2019; Qureshi et al., 2011). However, antigen-specific PTEN^KO^ Tregs showed no alteration in total CTLA-4 expression. It remains to be determined whether PTEN regulates CTLA-4 cycling to and from the cell surface, thought to be important for the transendocytic activity of CTLA-4 (Qureshi et al., 2011).

Curiously, the altered T cell- and APC-suppressive activity of PTEN^KO^ Tregs in vitro did not translate into a functional difference in vivo. We employed a xenogeneic model of GVHD, commonly used to validate human Treg performance (Dawson et al., 2019; Hippen et al., 2011). In this model, human CD8^+^ T cells but not CD4^+^ T cells are necessary and sufficient to induce disease (Søndergaard et al., 2013); non-T cells have limited engraftment and no known pathogenic role (van Rijn et al., 2003). T cell pathogenicity is dependent on recognition of mouse MHC class I/II and CD80/CD86 costimulation, though it is unclear which cell types provide these latter signals (King et al., 2009; Søndergaard et al., 2013; Zaitsu et al., 2017). Thus, this model primarily measures the direct effects of Tregs on T cell proliferation/cytokine production, potentially via CTLA-4 in tandem with other mechanisms. It is tempting to speculate that combined effect of both defective CTLA-4-dependent APC suppression and superior T cell suppression by PTEN^KO^ Tregs are represented in vivo, resulting in a net comparable PTEN^KO^ Treg performance to Cas9 Tregs.

Finally, other PI3K-AKT-targeting phosphatases may compensate for loss of PTEN function to maintain Treg stability and aspects of their suppressive capacity. In addition to PTEN, the AKT phosphatase PHLPP1 is co-recruited to the immunological synapse of Tregs upon TCR activation (Chen et al., 2017). PHLPP1 is required for mouse Treg development and function (Patterson et al., 2011), though its role in human Tregs is unclear. SHIP-1, a PI3K-targeting phosphatase, prevents human Tregs from reprogramming into Th1 cells (Lucca et al., 2019), and PP2A, which dephosphorylates AKT (T308), has been found to potentiate IL-2 signals in human Tregs (Ding et al., 2019; Sharabi et al., 2019). It is likely that multiple phosphatases operate in tandem to support different aspects of human Treg function.

Collectively, our findings suggest that, in contrast to previous studies, PTEN is responsible for a limited range of human Treg functions: PTEN is required for human Treg suppression of APC costimulation but is dispensable for their lineage stability and their ability to suppress T cell activity. Notably, defective PTEN^KO^ Treg suppression of APC costimulation was not revealed by two commonly used assays of Treg function, underscoring the importance of evaluating human Treg function from a variety of angles.

## Materials and Methods

### CRISPR editing reagents and quantification

CRISPR/Cas9 gRNA target sequences are in **Table S1**. crRNA and tracrRNA (200 μM each; both IDT) were duplexed (1:1 molar ratio) by incubation for 5 min at 95°C, followed by gradual cooling to room temperature. crRNA:tracrRNA duplex was complexed with SpCas9-2xNLS (40 μM; QB3 MacroLab) (2:1 molar ratio) by incubation for 10 min at room temperature.

To quantify indel formation after CRISPR editing, genomic DNA was isolated 3 or 8 days after editing by a QIAamp DNA Mini Kit or Micro Kit (QIAGEN). The genomic region flanking the CRISPR/Cas9 cleavage site was PCR-amplified, Sanger-sequenced, and chromatograms uploaded to the ICE (Synthego; https://ice.synthego.com/) web tool for decomposition (Hsiau et al., 2019). Primer sequences are in **Table S2**.

### Primary cell isolation and culture

Donor sex and age are in **Table S3**. For Treg isolation, CD4^+^ T cells were enriched by Lymphoprep and RosetteSep Human CD4^+^ T Cell Enrichment Cocktail (both STEMCELL Technologies). CD4-enriched cells were then CD25-enriched using CD25 MicroBeads II (Miltenyi Biotec), followed by flow sorting with a FACSAria IIu (BD Biosciences) or MoFlo Astrios (Beckman Coulter). Gating strategies (**Fig. S1A**): nTreg (CD4^+^CD25^hi^CD127^lo^CD45RA^+^), mTreg (CD4^+^CD25^hi^CD127^lo^CD45RA^−^), Tconv (CD4^+^CD25^lo^CD127^hi^). PBMCs were isolated by Lymphoprep. CD3^+^ T cells were isolated by Lymphoprep and RosetteSep Human T Cell Enrichment Cocktail. Monocytes were isolated from PBMCs using an EasySep Human Monocyte Isolation Kit (all STEMCELL Technologies).

Unless otherwise specified, T cells were cultured in X-VIVO 15 (Lonza) supplemented with 5% (v/v) human serum (WISENT), 1% (v/v) penicillin-streptomycin (Gibco), 2 mM GlutaMAX, (Gibco), and 15.97 mg/L phenol red (Sigma-Aldrich) at 37°C, 5% CO2. During the differentiation of moDCs, the medium above was additionally supplemented with 1 mM sodium pyruvate (STEMCELL Technologies).

### Treg expansion with CRISPR editing

Tregs and Tconvs were preactivated for 5 days with IL-2 (Proleukin, Novartis; 1000 IU/ml for Tregs, 100 IU/ml for Tconvs), anti-CD3 (OKT3, 0.1 μg/ml, University of British Columbia Antibody Lab), and 1:1 artificial APCs. Mouse L cells (ATCC CRL-2648; male) overexpressing hCD32, hCD80, hCD58 and gamma-irradiated (75 Gy) served as artificial APCs (de Waal Malefyt et al., 1993). Media and IL-2 were refreshed every 2–3 days.

Tregs were CRISPR-edited by electroporation with a Neon Transfection System 10 uL Kit (Invitrogen) as previously described (Lam et al., 2021): cells were washed twice with PBS, resuspended in Buffer T (≤20 × 10^6^ cells/ml), and electroporated at 1400 V / 30 ms / 1 pulse with Cas9 or CRISPR/Cas9 ribonucleoprotein (20 pmol Cas9 per transfection), then immediately transferred into prewarmed antibiotic-free medium and expanded for 7 days with IL-2, anti-CD3, and 1:1 artificial APCs as above (total 12 days). Tconvs (not electroporated) were preactivated and expanded in parallel. See **Fig. S1B** for timeline.

Unless otherwise indicated, for all assays, cells were rested overnight with reduced IL-2 as above (total 13-day culture). For **Fig. S2**, 12 day-expanded cells were further expanded for 7 days with ImmunoCult Human CD3/CD28/CD2 T Cell Activator (25 μl/ml; STEMCELL Technologies), then rested overnight with reduced IL-2 as above (total 20-day culture). See **Fig. S1B** for timeline.

To generate gene-edited A2-CAR Tregs, nTregs from HLA-A2^−^ donors were preactivated, electroporated, and expanded as above (total 13-day culture). In addition, 1 day after preactivation, cells were transduced with a bidirectional-promoter lentiviral vector encoding EF1α-driven A2CAR-CD28-CD3? and minimal CMV-driven truncated nerve growth factor receptor (ΔNGFR), referred to as A2-CAR (multiplicity of infection = 10) (MacDonald et al., 2016). After expansion, NGFR^+^ cells were bead-purified by an EasySep Human CD271 Positive Selection Kit II (STEMCELL Technologies). See **Fig. S4A** for timeline.

### Seahorse extracellular flux analysis

Tregs and Tconvs were assayed immediately after total 12-day expansion. As previously described (van der Windt et al., 2016), cells were resuspended in unbuffered RPMI 1640 (Sigma-Aldrich) supplemented with D-glucose (10 mM; Sigma-Aldrich), glutamine (4 mM), and sodium pyruvate (1 mM; Gibco). Cells were plated in triplicate (200,000 cells/well) in a Seahorse XFe96 assay plate (Agilent Technologies) coated with poly-D-lysine (50 μg/ml; Sigma-Aldrich). Extracellular acidification and oxygen consumption rates were measured using the mitochondrial stress test procedure with a Seahorse XFe96 Analyzer (Agilent Technologies) under basal conditions and upon the sequential addition of oligomycin (1 μM), FCCP (1.5 μM), and rotenone (100 mM) + antimycin A (1 mM; all Sigma-Aldrich).

### Cytokine detection

Secreted cytokines were assessed with LEGENDplex Human Th Panel 12-plex and analysed with Qognit software (both BioLegend). For cytokine secretion by Tregs, cells were activated with 100 IU/ml IL-2 and 1:1 Dynabeads Human T-Expander CD3/CD28, and culture supernatants were collected after 4 days. For suppression of cytokine secretion, supernatants from 4-day Treg:CD3^+^ T cell co-cultures were collected. Percent suppression of cytokine secretion was calculated as: (1 – (pg/ml cytokine of sample / pg/ml cytokine of positive control)) × 100%. For intracellular cytokine expression, Tregs were activated with PMA (10 ng/ml), ionomycin (500 ng/ml), and brefeldin A (10 μg/ml; all Sigma-Aldrich) for 6 h at 37°C.

### FOXP3 TSDR methylation

Genomic DNA was isolated and bisulfite-converted with an EZ DNA Methylation-Direct Kit (Zymo), PCR-amplified, and pyrosequencing performed with a PyroMark Q96 MD (QIAGEN). Primer sequences are in **Table S3**.

### In vitro suppression of T cell proliferation

Tregs and allogeneic CD3^+^ T cells were labelled with Cell Proliferation Dye eFluor 670 and eFluor 450 (Invitrogen), respectively, then co-cultured at the indicated ratios and activated with Dynabeads Human T-Expander (1 bead:16 CD3^+^ T cells; Gibco) for 4 days. CD3^+^ T cells activated alone served as positive controls. In all cases, CD3^+^ T cells were plated at 50,000 cells/well in a 96-well U-bottom plate. Percent suppression of CD4^+^ T cell and CD8^+^ T cell proliferation was calculated using division index: (1 – (division index of sample / division index of positive control)) × 100%.

### In vitro suppression of dendritic cells

HLA-A2^+^ monocytes were differentiated into moDCs for 7 days with GM-CSF (50 ng/ml) and IL-4 (100 ng/ml; both STEMCELL Technologies); media and cytokines were refreshed every 2–3 days. Maturation was induced by a combination of IL-1β (10 ng/ml; Invitrogen), IL-6 (100 ng/ml, Invitrogen), TNF-α (10 ng/ml; Invitrogen), PGE2 (1 μg/ml; Tocris) (all last 2 days), and IFN-γ (50 ng/ml, Invitrogen) (last 1 day) (**Fig. S4A**). Maturation was confirmed by high expression of CD80, CD83, CD86, and HLA-DR. Suppression of the costimulatory potential of moDCs was evaluated as previously described (Huang et al., 2021): A2-CAR Tregs (250,000/well) and mature HLA-A2^+^ moDCs (50,000/well) were co-cultured (5 Treg:1 moDC) with IL-2 (50 IU/ml) for 4 days in a 96-well U-bottom plate. Immature or mature moDCs cultured alone served as controls. Expression of CD80, CD83, CD86, and HLA-DR by CD11c^+^ moDCs after co-culture were determined by flow cytometry.

### In vivo suppression of xenogeneic graft-versus-host disease

Cas9 or PTEN^KO^ nTregs from HLA-A2^+^ donors were expanded for 12 days as above, then cultured overnight with 1000 IU/ml IL-2 (total 13-day culture), then resuspended in PBS (**Fig. S3A**). Allogeneic HLA-A2^−^ PBMCs were thawed, incubated with DNase I (10 μg/ml; STEMCELL Technologies) for 10 min at room temperature, washed with PBS, and resuspended in PBS. 6- to 10-week-old female NSG nice (NOD.Cg-*Prkdc*^*scid*^*Il2rg*^*tm1Wjl*^/SzJ, Jackson Laboratory; bred in-house) received total body X-ray irradiation (150 cGy) 1 day prior to intravenous injection of PBMCs (8 × 10^6^/mouse) and expanded nTregs (4 × 10^6^/mouse or 2 × 10^6^/mouse). The degree of GVHD was assessed thrice weekly based on weight loss, activity, posture, fur texture and skin integrity, and pain with a score of 0 to 3 (Cooke et al., 1996). Mice with a score of 3 in any category or a total score of 6 were euthanized, in which case a value of 6 was assigned. Blood from saphenous vein was collected once weekly for immune monitoring.

The same PBMC donor was used for all mice. PBMC-only and PBS-injected mice were included in all cohorts. Mice from different experimental groups were co-housed.

### Western blotting

Cells were resuspended in PBS, lysed with an equal volume of 2X Laemmli buffer (2.2% SDS, 10% glycerol, 0.1 M Tris, pH 6.8) for 5 min at 95°C, sonicated for 4 min, and quantified with a Pierce BCA Protein Assay Kit (Thermo Scientific). 15 μg protein was resolved in a 10% SDS gel, transferred to a polyvinylidene difluoride membrane, and blocked with 5% (w/v) bovine serum albumin (Sigma-Aldrich). Membranes were subsequently probed with the antibodies in **Table S4** in 5% (w/v) bovine serum albumin and detected by chemiluminescence (Pierce ECL Western Blotting Substrate, Thermo Scientific).

### Flow cytometry

Flow cytometry was performed in adherence to “Guidelines for the use of flow cytometry and cell sorting in immunological studies (third edition)” (Cossarizza et al., 2021). Flow cytometric antibodies are in **Table S5**. Cells were stained with Fixable Viability Dye eFluor 780 (Invitrogen) and surface proteins in PBS (Gibco) or 1X Brilliant Stain Buffer (BD Biosciences) for 20 min at room temperature before fixation/permeabilization with the eBioscience Foxp3 / Transcription Factor Staining Buffer Set (Invitrogen) and intracellular staining for 40 min at room temperature. Data were acquired with a FACSymphony A5, LSRII (both BD Biosciences), or CytoFLEX (Beckman Coulter) and analysed with FlowJo software (v10.7.1; BD). Total CTLA-4 expression was determined by staining after fixation and permeabilization.

For phosphoflow, 12 day-expanded cells were rested overnight with reduced IL-2 (100 IU/ml for Tregs, none for Tregs), then starved overnight without serum or IL-2 (total 14-day culture). In some cases as indicated, cells were preincubated with anti-CD3 (OKT3, 1 μg/ml) and anti-CD28 (CD8.2, μg/ml; BD Biosciences) for 15 min on ice, washed, then cross-linked with AffiniPure F(ab′)_2_ Fragment Goat Anti-Mouse IgG (20 μg/ml; Jackson ImmunoResearch) (“TCR”) or without (“Unstim”) for 5 min at 37°C. Cells were immediately fixed with Cytofix Fixation Buffer for 15 min at 37°C, washed, permeabilized with Perm Buffer III for 30 min at 4°C (both BD Biosciences), washed twice, and stained for phospho-proteins in PBS for 40 min at room temperature.

For in vivo studies, blood and physically dissociated spleen samples were lysed of red blood cells by incubation with 1X eBioscience RBC Lysis Buffer (Invitrogen) for 3min (blood) or 2×3 min (spleen) at room temperature, washed twice with PBS, and preincubated with Mouse BD Fc Block (BD Biosciences; clone 2.4G2) for 10 min at room temperature before staining for surface and intracellular antigens as above. 123count eBeads Counting Beads (Invitrogen) were added immediately before acquisition to obtain absolute counts.

All live single cells were first gated as CD4^+^ (**Fig. S1A, top row**), except for suppression assays for T cell proliferation (CD4^+^ or CD8^+^) (**Fig. S5A**), moDCs (CD11c^+^) (**Fig. S5B**), and xenogeneic graft-versus-host disease (as indicated) (**Fig. S5C**).

### Statistical analysis

Normality was assumed. The n and p values are indicated in the figures or figure legends (p < 0.05 was considered statistically significant). Significance between two groups was determined by paired t-test. For comparison of more than two groups, matched 1-way or 2-way ANOVA with Dunnett’s or Sidak’s multiple comparisons test was used as appropriate. For suppression of T cell proliferation and cytokine secretion, t-test of the areas under the curve was used, considering all degrees of freedom, as previously described (Akimova et al., 2016). Analysis was performed with Prism software (v9.0.1; GraphPad). For pairwise comparisons of survival curves, log-rank test was used.

## Abbreviations

APC: antigen-presenting cell
CAR: chimeric antigen receptor
gRNA: guide RNA
GVHD: graft-versus-host disease
mTreg: memory regulatory T cell
moDC: monocyte-derived dendritic cell
nTreg: naïve regulatory T cell
Tconv: CD4^+^ conventional T cell
Treg: regulatory T cell
TSDR: Treg-specific demethylation region

## Data Availability Statement

The data that support the findings of this study are available from the corresponding author upon reasonable request.

## Ethics Approval Statement

Collection of human samples (buffy coats or leukapheresis products) was performed following written informed consent and in accordance with protocols approved by the UBC Clinical Research Ethics Board and Canadian Blood Services Research Ethics Board, or purchased from STEMCELL Technologies. Animal protocols were approved by the UBC Animal Care Committee.

## Author Contributions

Conceptualization: AJL, MKL

Methodology: AJL, KAWH, JKG, MM, RIKG

Investigation: AJL, MH, PU, CMW, JKG, MS, MM

Formal Analysis: AJL, MH, KAWH, JKG

Writing—Original Draft: AJL, MKL

Writing—Review and Editing: AJL, MH, KAWH, PU, CMW, JKG, MS, MM, RIKG, MKL

Supervision: MM, RIKG, MKL

Funding Acquisition: MKL

## Acknowledgements

We thank the UBC Nucleic Acid Protein Service and the BC Children’s Hospital Research Institute (BCCHR) DNA Sequencing and Flow Core Facilities for DNA sequencing, instrumentation, and assistance with flow sorting. This work was supported by a grant from the Canadian Institutes of Health Research (CIHR; FDN-154304 to MKL). AJL is supported by a CIHR doctoral research award, KAWH is supported by fellowships from CIHR and the Juvenile Diabetes Research Foundation, CMW is supported by a BCCHR graduate studentship, and MKL receives a BCCHR salary award and holds a Tier 1 Canada Research Chair in Engineered Immune Tolerance.

