## Supplemental Figures & Tables for "PTEN is required for human Treg suppression of costimulation"

Fig. S1. Flow sorting and expansion timeline for CRISPR-edited Tregs.

Fig. S2. Human Tregs retain lineage stability and superior T cell-suppressive capacity with prolonged $PTEN^{KO}$.

Fig. S3. Extended timeline of xenogeneic GVHD model and phenotype of transferred Tregs.

Fig. S4. Expansion timeline, phenotype, and T cell-suppressive capacity of human A2-CAR $PTEN^{KO}$ Tregs.

Fig. S5. Gating strategies for in vitro and in vivo Treg suppression assays.

Table S1. CRISPR gRNA target sequences.

Table S2. Primer sequences for PCR amplification and sequencing.

Table S3. Donor age and sex, if known.

Table S4. Antibodies for western blotting.

Table S5. Antibodies for flow cytometry.

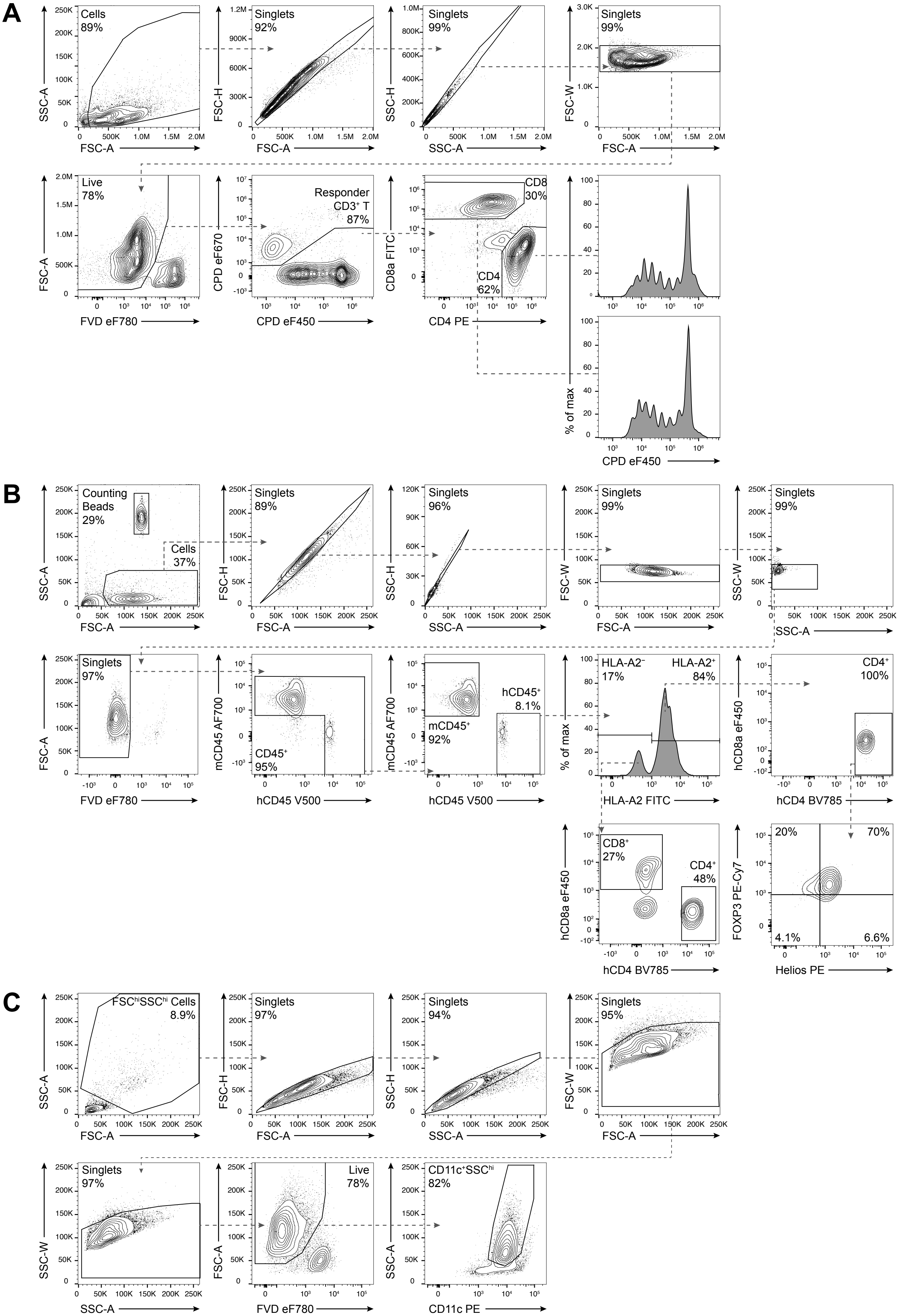


**Fig. S5. Gating strategies for in vitro and in vivo Treg suppression assays.**
(**A**) Identification of CD4⁺ and CD8⁺ T cells (Cell Proliferation Dye (CPD) eF450-labelled) and Tregs (CPD eF670-labelled) after Treg:CD3⁺ T cell co-culture. (**B**) Identification of PBMCs (hCD45⁺HLA-A2⁻) and Tregs (hCD45⁺HLA-A2⁺CD4⁺) in mouse blood from a xenogeneic GVHD model. (**C**) Identification of moDCs (CD11c⁺) after co-culture with Tregs.

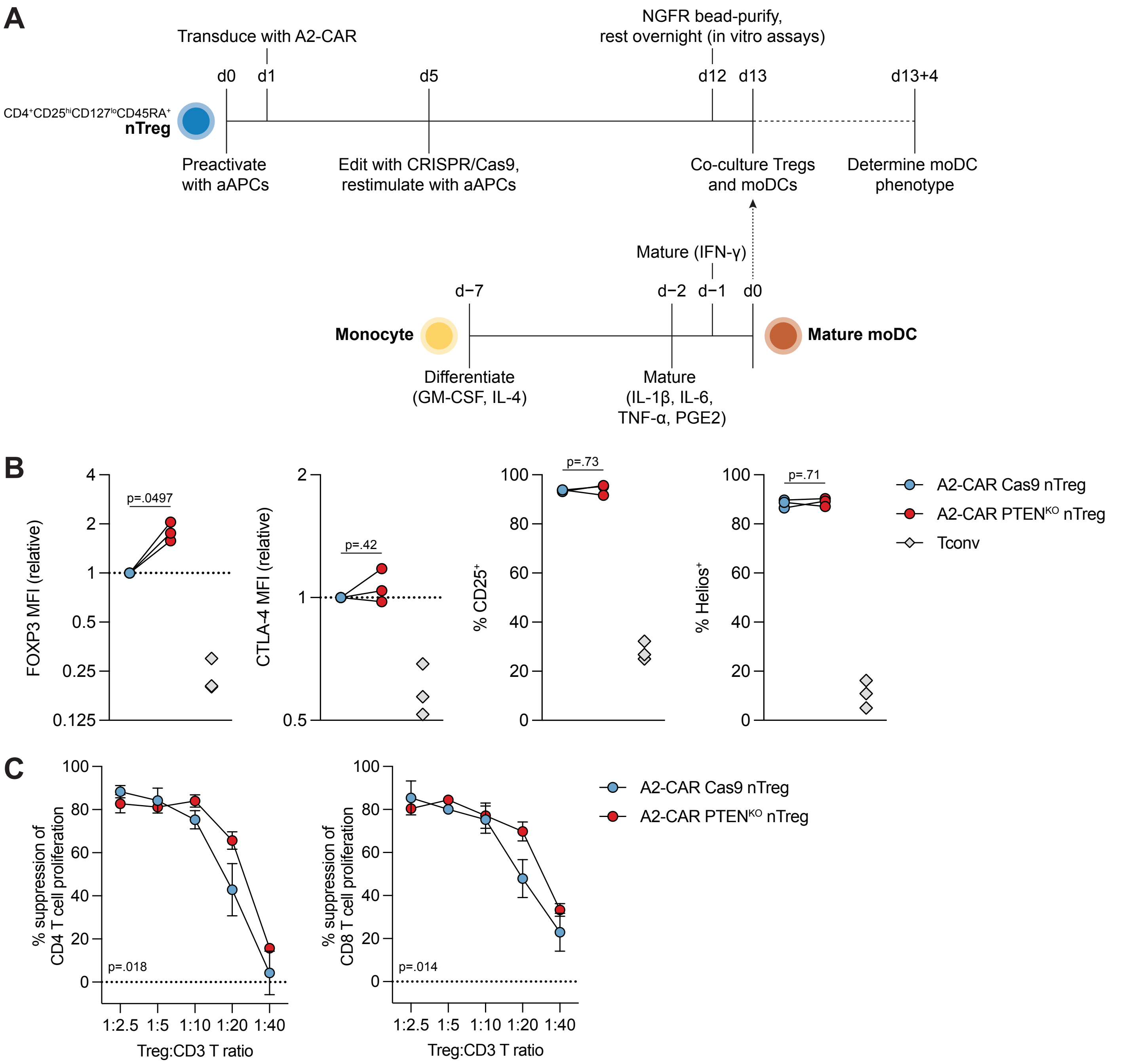


**Fig. S4. Expansion timeline, phenotype, and T cell-suppressive capacity of human A2-CAR PTEN$^{KO}$ Tregs.**
(**A**) To generate gene-edited A2-CAR nTregs to assess their suppressive capacity of moDCs, sorted nTregs were transduced, edited, and expanded as indicated. moDCs were differentiated in parallel as indicated. (**B**) Expression of FOXP3, CTLA-4, CD25, and Helios by flow cytometry (n=3, 1 experiment). (**C**) nTregs were co cultured with allogeneic CD3+ T cells and TCR-activated (4 days). Suppression of CD4$^{+}$ T cell (left) and CD8$^{+}$ T cell (right) proliferation (n=2, 1 experiment). Dots in (**B**) represent individual donors. (**C**) depicts mean±SEM. Significance determined by paired t-test in (**B**) and t-test of the areas under the curve in (**C**). Tconvs shown for reference. MFI, geometric mean fluorescence intensity.

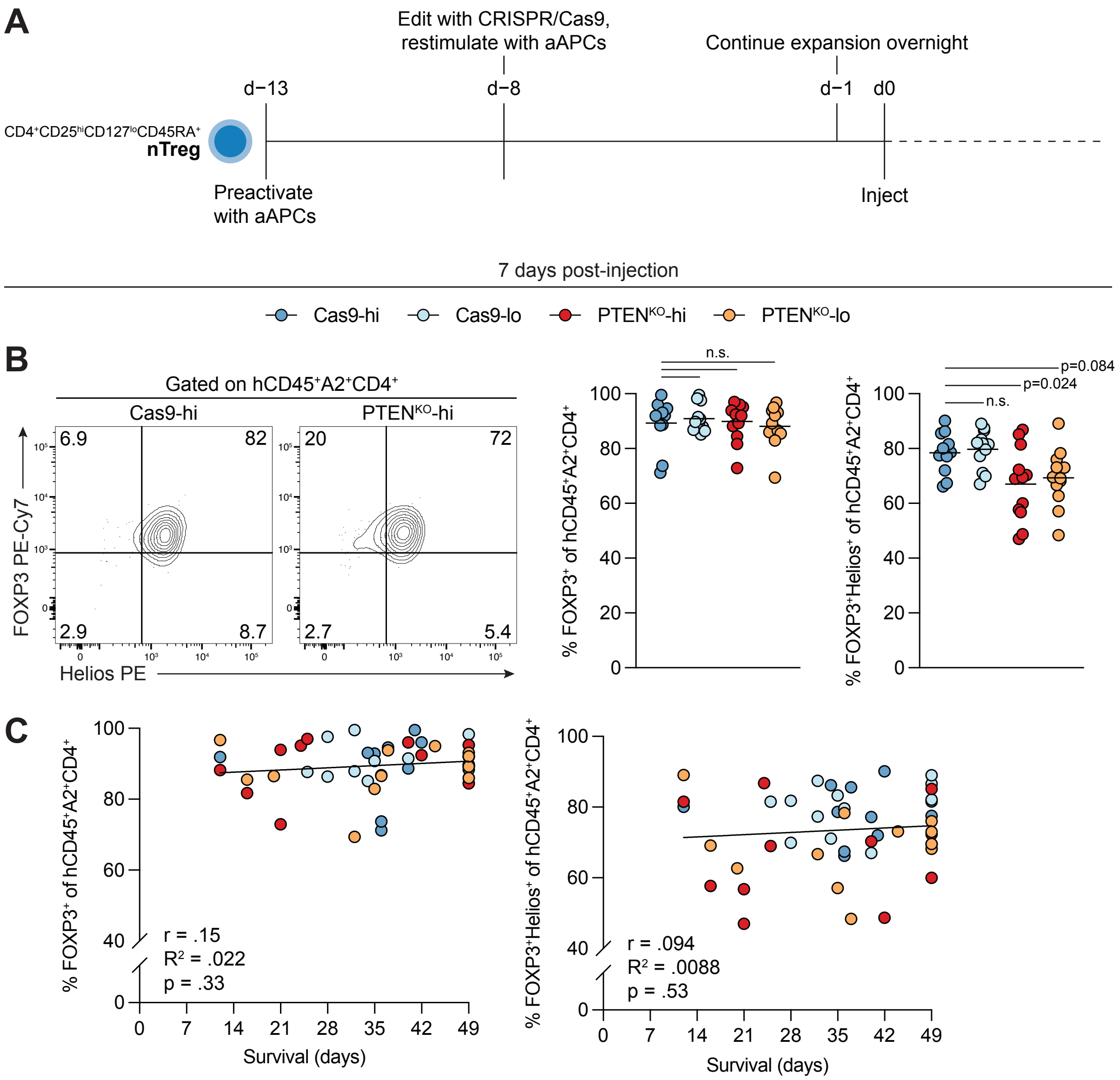


**Fig. S3. Extended timeline of xenogeneic GVHD model and phenotype of transferred Tregs.**
(**A**) Sorted nTregs were edited, expanded, and injected as indicated. See Fig. 5A for in vivo timeline after injection (day 0 onwards). (**B**) Example (left) and quantified (right) expression of FOXP3 and Helios in transferred Cas9 and PTEN$^{KO}$ nTregs, 7 days post-injection. (**C**) Correlation of FOXP3$^+$ or FOXP3$^+$Helios$^+$ expression in transferred Tregs (day 7) with mouse survival time. Dots in (**B–C**) represent individual mice. Lines in (**C**) represent lines of best fit. Significance in (**B**) determined by 1-way ANOVA with Dunnett's multiple comparisons test (all groups compared to Cas9-hi). Pearson correlation coefficient (r) shown in (**C**).

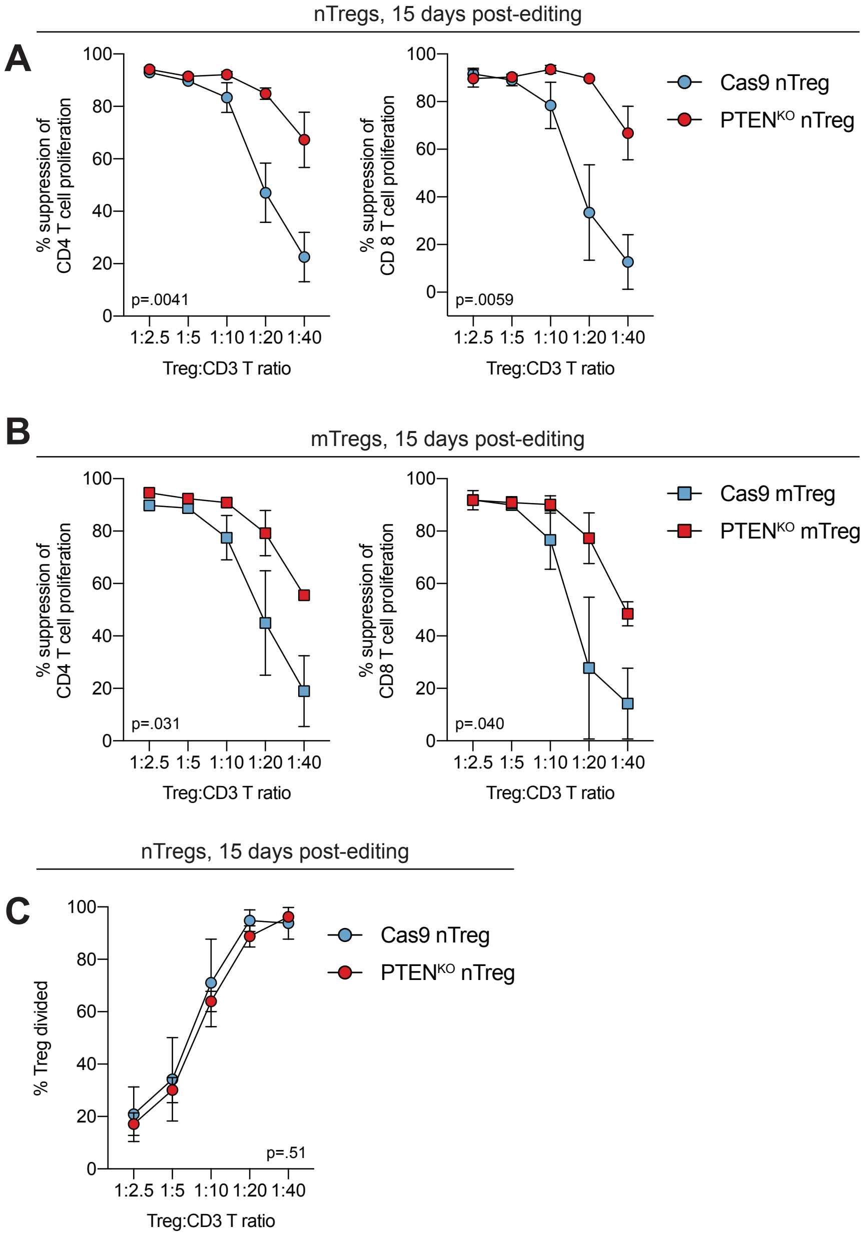


**Fig. S2. Human Tregs retain lineage stability and superior T cell-suppressive capacity with prolonged $PTEN^{KO}$.** (**A–C**) Cas9 and $PTEN^{KO}$ nTregs (**A, C**) and mTregs (**B**) were expanded for total 20 days, then co-cultured with allogeneic $CD3^+$ T cells at the indicated ratios and TCR-activated (4 days). (**A**) nTreg-mediated suppression of $CD4^+$ T cell (left) and $CD8^+$ T cell (right) proliferation (n=3, 2 experiments). (**B**) mTreg-mediated suppression of CD4⁺ T cell (left) and $CD8^+$ T cell (right) proliferation (n=2, 2 experiments). (**B**) Proliferation of nTregs in the nTreg:$CD3^+$ T cell co-culture (n=3, 2 experiments). (**A–C**) depicts mean±SEM. Significance in (**A–C**) determined by t-test of the areas under the curve. Tconvs shown for reference. MFI, geometric mean fluorescence intensity.

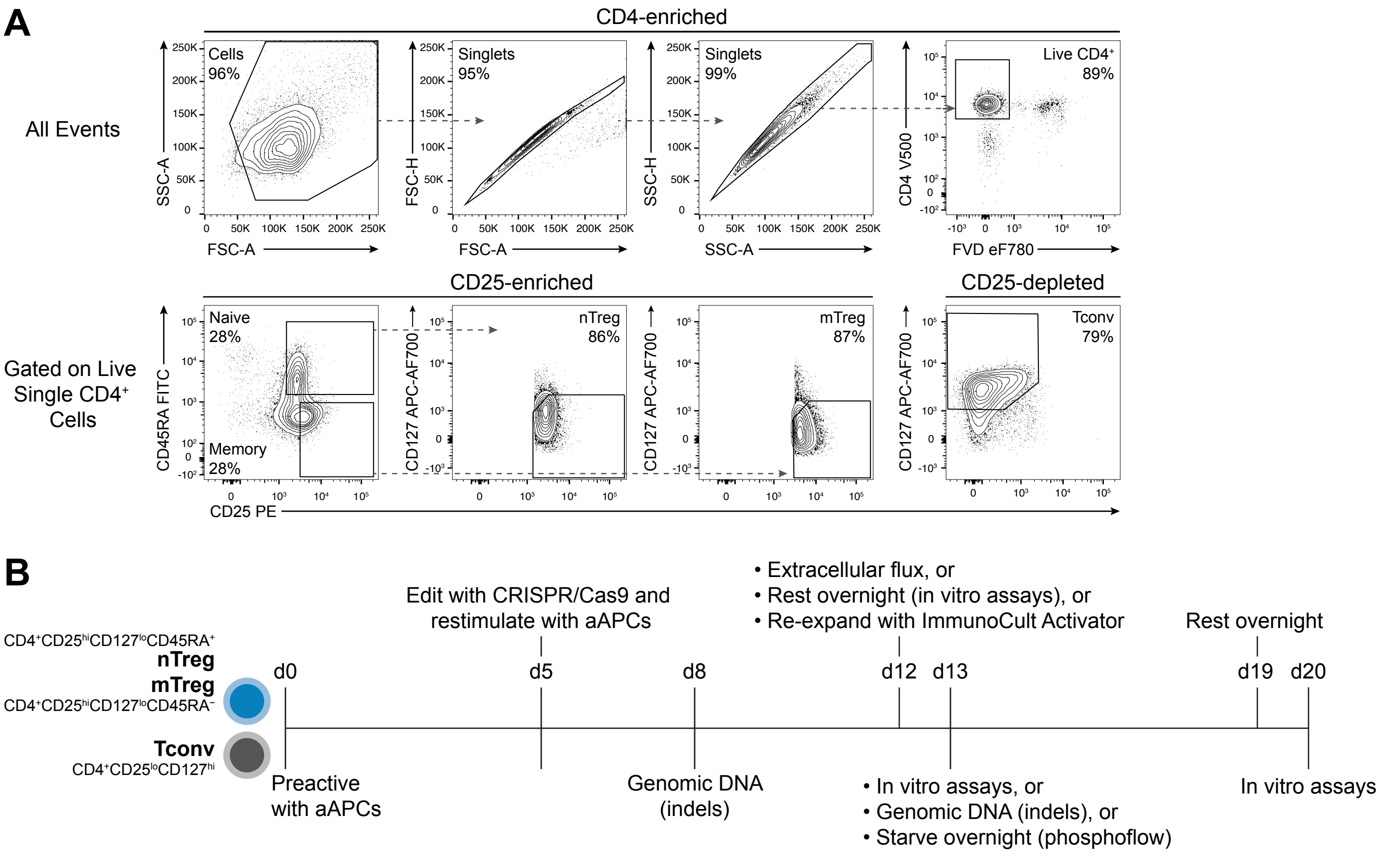


**Fig. S1. Flow sorting and expansion timeline for CRISPR-edited Tregs.**
(**A**) Flow sorting strategy for naive Tregs (nTreg; CD4$^{+}$CD25$^{hi}$CD127$^{lo}$CD45RA$^{+}$), memory Tregs (mTreg; CD4$^{+}$CD25$^{hi}$CD127$^{lo}$CD45RA$^{-}$), and conventional T cells (Tconv; CD4$^{+}$CD25$^{lo}$CD127$^{hi}$). (**B**) For in vitro studies, sorted cells were edited, expanded, and assayed as indicated. aAPC, artificial antigen-presenting cell; FVD, Fixable Viability Dye.

**Table S1. CRISPR gRNA target sequences.**

| gRNA | Target sequence (5′–3′) |
|---|---|
| PTEN CR1 | TTATCCAAACATTATTGCTA |
| PTEN CR2 | TATCCAAACATTATTGCTAT |
| PTEN CR3 | ACAGATTGTATATCTTGTAA |
| PTEN CR4 | ACCTCTGCAATTAAATTTGG |
| PTEN CR5 | ACCGCCAAATTTAATTGCAG |
| PTEN CR6 | TTGATGATGGCTGTCATGTC |
| PTEN CR7 | CCTACCTCTGCAATTAAATT |
| PTEN CR8 | CCAAATTTAATTGCAGAGGT |
| PTEN CR9 | GACTGGGAATAGTTACTCCC |

**Table S2. Primer sequences for PCR amplification and sequencing.**

| Product | Forward (5′–3′) | Reverse (5′–3′) | Sequencing (5′–3′) |
|---|---|---|---|
| PTEN CR1, CR2 | TCTTTTCAGGCAGGTGTCAA | TGGTGACCAGCATTTTATGG | CCGTGAGTTTCTGTTTTTCTCA |
| PTEN CR3 | TGATGGGAAAATGATGTCTGA | CATGAATCTGTGCCAACAATG | Reverse |
| PTEN CR4, CR5, CR7, CR8 | TGGCATCACAAGTTTTTAAGCA | TGCTGCACTTTAGTCTTCCTGA | Reverse |
| PTEN CR6 | GCAGCCGTTCGGAGGATTA | AGGCAAGAGTTCCGTCTAGC | Reverse |
| PTEN CR9 | CAGAGCGCTGTTGTGACCTT | TGGAAGGATGAGAATTTCAAGCACT | Forward |
| *FOXP3* TSDR | AGAAATTTGTGGGGTGGGGTAT | ATCTACATCTAAACCCTATTATCACAACC (biotinylated) | AGAAATTTGTGGGGTGGG |

**Table S3. Donor age and sex, if known.**

| Donor ID | Tissue Type | Sex | Age |
|---|---|---|---|
| 1473 | PB | F | N/A |
| 0452 | PB | M | 23 |
| 0482 | PB | M | 42 |
| 1924 | PB | F | 42 |
| 7608 | PB | F | 17 |
| 3142 | PB | F | 42 |
| 0807 | PB | M | 20 |
| 5100 | PB | M | N/A |
| 8705 | PB | M | N/A |
| 3694 | PB | F | N/A |
| 1715 | PB | M | N/A |
| 7852 | PB | F | 63 |
| 0355 | PB | F | 23 |
| 0358 | PB | M | 26 |
| 7579 | PB | M | 26 |
| 7564 | PB | F | 32 |
| 7549 | PB | M | 38 |
| D3565 | PB | M | 38 |
| 3392 | PB | M | 34 |
| 6087 | PB | M | 29 |
| 2538 | PB | M | 34 |
| 3984 | PB | F | N/A |

**Table S4. Antibodies for western blotting.**

| Target | Clone | Host | Dilution | Company | Cat# |
|---|---|---|---|---|---|
| p-AKT (S473) | 193H12 | Rabbit | 1/1000 | Cell Signaling Technology | 4058 |
| p-FOXO1 (S256) | Polyclonal | Rabbit | 1/1000 | Cell Signaling Technology | 9461 |
| AKT | 40D4 | Mouse | 1/1000 | Cell Signaling Technology | 2920 |
| Beta-actin | Polyclonal | Rabbit | 1/1000 | Cell Signaling Technology | 4967 |
| FOXO1 | C29H4 | Rabbit | 1/1000 | Cell Signaling Technology | 2880 |
| PTEN | D4.3 | Rabbit | 1/1000 | Cell Signaling Technology | 9188 |
| Mouse IgG, HRP-linked | Polyclonal | Goat | 1/5000 | Sigma-Aldrich | AP127P |
| Rabbit IgG, HRP-linked | Polyclonal | Goat | 1/1000 | Cell Signaling Technology | 7074 |

**Table S5. Antibodies for flow cytometry.**

| Target | Clone | Fluorophore | Company |
|---|---|---|---|
| CD4 | OKT4 | PE | Invitrogen |
| CD4 | RPA-T4 | V500 | BD Biosciences |
| CD4 | SK3 | BUV395 | BD Biosciences |
| CD4 | SK3 | BUV496 | BD Biosciences |
| CD4 | SK3 | BV480 | BD Biosciences |
| CD4 | SK3 | BV785 | BD Biosciences |
| CD8a | HIT8a | FITC | Invitrogen |
| CD8a | SK1 | eFluor 450 | Invitrogen |
| CD11c | B-ly6 | PE | BD Biosciences |
| CD25 | 4E3 | PE | Miltenyi Biotec |
| CD25 | 2A3 | BB515 | BD Biosciences |
| CD25 | 2A3 | BUV395 | BD Biosciences |
| CD45 (mouse) | 30-F11 | AF700 | Invitrogen |
| CD45 | HI30 | V500 | BD Biosciences |
| CD45RA | HI100 | FITC | Invitrogen |
| CD80 | L307.4 | BB515 | BD Biosciences |
| CD83 | HB15e | BV421 | BioLegend |
| CD86 | 5C3 | PE-Cy7 | Invitrogen |
| CD127 | eBioRDR5 | eFluor 450 | Invitrogen |
| CD127 | R34.34 | APC-AF700 | Beckman Coulter |
| CD152 (CTLA-4) | BNI3 | APC | BD Biosciences |
| CD152 (CTLA-4) | BNI3 | BUV496 | BD Biosciences |
| CD152 (CTLA-4) | BNI3 | BV785 | BD Biosciences |
| FOXP3 | 236A/E7 | PE | Invitrogen |
| FOXP3 | 236A/E7 | PE-Cy7 | Invitrogen |
| Helios | 22F6 | AF647 | BioLegend |
| Helios | 22F6 | PE | BioLegend |
| HLA-A2 | BB7.2 | FITC | BD Biosciences |
| HLA-A2 | BB7.2 | PE | BD Biosciences |
| HLA-A3 | GAP.A3 | FITC | Invitrogen |
| HLA-DR | G46-6 | BV480 | BD Biosciences |
| IFN-γ | B27 | BUV395 | BD Biosciences |
| IL-10 | JES3-9D7 | BV421 | BioLegend |
| IL-17A | N49-653 | BV785 | BD Biosciences |
| IL-2 | MQ1-17H12 | BUV737 | BD Biosciences |
| Myc | 9E10 | AF647 | UBC Antibody Lab |
| p-AKT (S473) | M89-61 | AF647 | BD Biosciences |
| p-AKT (T308) | J1-223.371 | PE | BD Biosciences |
| p-S6 (S235/S236) | cupk43k | eFluor 450 | Invitrogen |
